# Comparative microbiomes of three species of laboratory-reared *Periplaneta* cockroaches

**DOI:** 10.1101/824524

**Authors:** Seogwon Lee, Ju Yeong Kim, Myung-hee Yi, In-Yong Lee, Tai-Soon Yong

**Author notes:** Address correspondence to Tai-Soon Yong.

## Abstract

Cockroaches are the most primitive insects, and are often used as the basic insect model in many studies. Three species of *Periplaneta* cockroaches were raised in the laboratory for many generations under the same conditions. We conducted 16S rRNA-targeted high-throughput sequencing to evaluate the overall bacterial composition in the microbiomes of three species of cockroaches. The number of operational taxonomic units (OTUs) was not significantly different between the three cockroach species. With respect to the Shannon and Pielou indexes, the microbiome of *Periplaneta americana* presented higher values than that of either *P. japonica* or *P. fulginosa*. In terms of species composition, endosymbionts accounted for over half of all the bacterial species in *P. japonica* and *P. fulginosa*. The beta diversity analysis showed that *P. japonica* and *P. fulginosa* exhibit a similar microbiome composition, which is different from that of *P. americana*. However, we also identified that *P. japonica* and *P. fulginosa* are hosts to distinct bacterial species. Thus, although the composition of the microbiome may vary based on multiple conditions, it is possible to identify distinct compositions of the microbiome among the different *Periplaneta* cockroach species even when individuals are reared under the same conditions.

**Importance:** Cockroaches inhabit various habitats—which are known to be related to their microbiome—and exhibit different features depending on the species. It is expected that their microbiomes would vary according to species depending on these features. Cockroach microbiomes are known to vary based on the diet or environmental shifts. In this study, we compare the diversity of bacteria in the three cockroach species under conditions of reduced diet and environment shifts. This study can establish the basic microbiomes of three *Periplaneta* species and can be the basic data for cockroach research.

## Introduction

Cockroaches are among the most primitive insects. They are often used as insect models to study reproductive, genetic, and insecticide-resistance mechanisms. Cockroaches are tolerant to both heat and cold, although this trait varies between species. For example, *Blatta materialis*, can tolerate temperatures of up to 48.1 °C (1), and *B. auricularis* has been successfully maintained at 10 °C for 14 days (2). Cockroaches are able to maintain a stable internal environment by regulating their water balance. These characteristics have allowed cockroaches to survive in various habitats, including tropical forests, deserts, and coastal areas (3). This habitat diversity has been found to be associated with the microbiomes of the cockroaches themselves (4).

Insect microbiomes affect nutrient recycling, provide protection from parasites and pathogens, and modulate immune responses. Cockroach microbiomes consist of horizontally transmitted microbes and vertically transmitted symbionts. The diversity of these microbiomes can vary depending on developmental stage, diet, and rearing practices (4). In a recent study, laboratory-reared and field-collected *Blattella germanica* presented distinct microbiomes, although they shared the same core bacterial taxa, which appear to differ depending on the location and diet (5). However, no significant microbiome differences have been observed in *Periplaneta americana* due to changes in diet, although this species has been found to present microbiome differences due to environmental factors (6).

Forty-seven species are included in the *Periplaneta* genus (7), and we have three species of *Periplaneta* genus in our laboratory. *Periplaneta americana* originated in Africa and is very common worldwide (7). This species measures about 4 cm in length (7) and is often found in commercial buildings (8). *Periplaneta fuliginosa* is another species of African origin and measures 3 cm in length. This species is widely distributed across the southeastern United States and Japan (9). *Periplaneta japonica*, which measures 2.5 cm in length, originated in Japan and is freeze tolerant (10). As previously stated, these features are known to be associated with the microbiomes of the cockroaches (4).

Since the features of each species are different within the *Periplaneta* genus, we expected that there would also be differences among *Periplaneta* microbiomes. As such, we conducted research to establish a microbiome that minimized the aforementioned differences that may have been due to diet and environmental factors. The laboratory-reared cockroaches used in this study were reared for many generations under the same laboratory conditions to minimize the differences brought about by diet and environmental factors. Then, we analyzed the cockroach microbiomes using 16S rRNA targeted high-throughput sequencing to compare the three cockroach species.

## Results

The average number of read counts assigned to *P. americana, P. japonica*, and *P. fuliginosa* were 49905 reads corresponding to 897 species (operational taxonomic units, OTUs), 56565 reads corresponding to 955 species, and 58013 reads corresponding to 878 species, respectively (Data Set S1). The rarefaction curve of all samples formed a plateau (Fig. S1). The number of OTUs was not significantly different between the three cockroach species (Fig. 1A). There were no significant differences in the phylogenetic index between species, although it was higher for *P. americana* than that for either *P. japonica* or *P. fulginosa* (Fig. 1B, p = 0.055, 0.262). In contrast, the Pielou and Shannon index values were significantly higher for *P. americana* than that for either *P. japonica* or *P. fulginosa* (Fig. 1C, p = 0.004).

**Figure 1.**
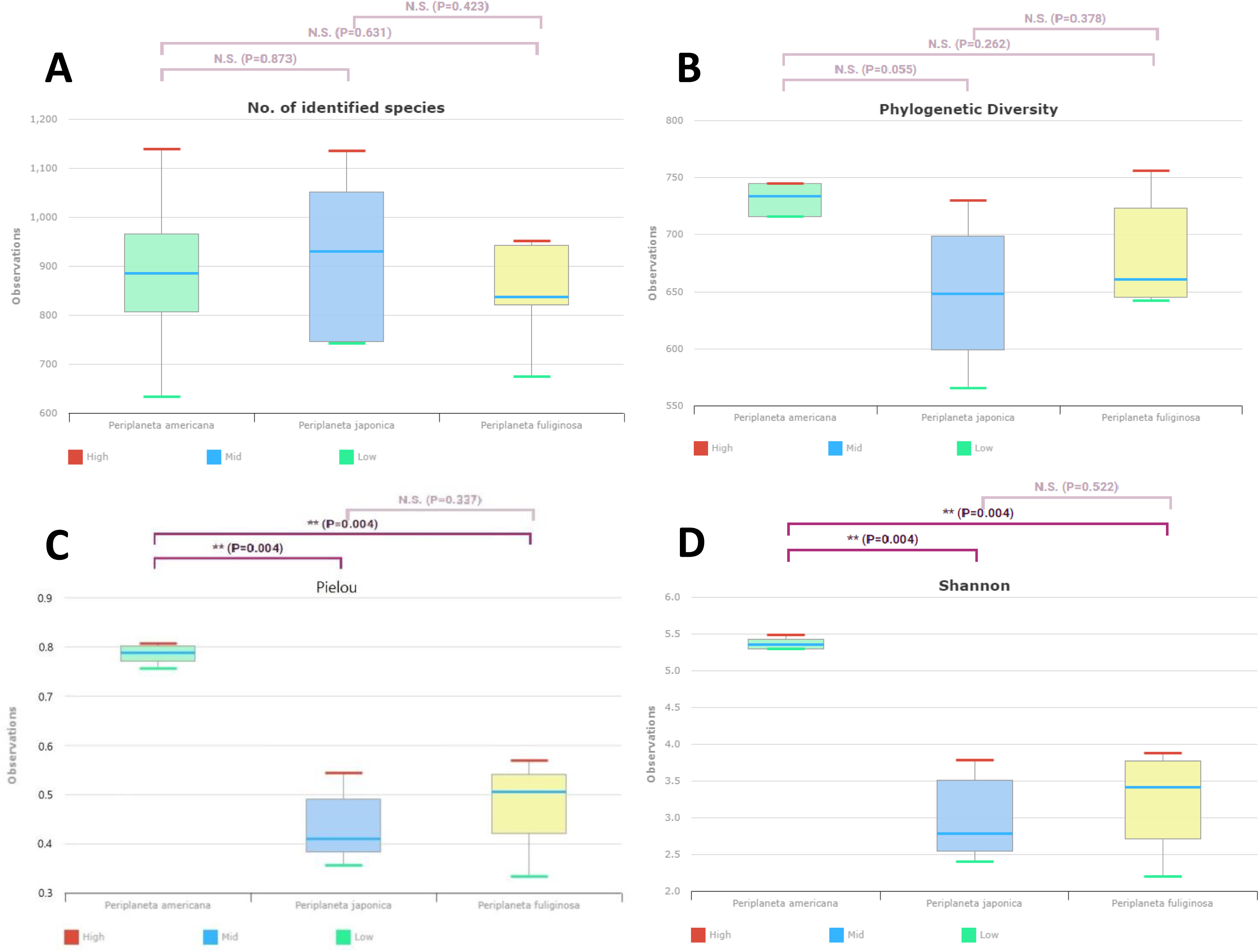
Box plots showing the alpha diversity (measurement of species richness, abundance, and equity within a habitat unit) of **(A)** the number of operational taxonomic units (OTUs) found in microbiome taxonomic profiling (MTP), **(B)** phylogenetic diversity (abundance), **(C)** Pielou diversity (equity), and **(D)** Shannon diversity (measurement of abundance and equity of the distribution of species) among the cockroach samples. Bars indicate the median, and the hinges represent the lower and upper quartiles. In panels **(A)** and **(B)** no statistically significant differences were observed between the three species of cockroaches. However, in panels **(C)** and **(D)**, statistically significant differences between *Periplaneta americana*, and *P. japonica* and *P. fuliginosa* were observed.

The UPGMA cluster analysis showed that the cockroaches were organized according to species; *P. japonica* and *P. fulginosa* clustered earlier and then joined *P. americana* (Fig. 2A). The results of the PCoA showed that even though all three groups clustered together, the *P. americana* samples were more tightly clustered than either the *P. japonica* or *P. fulginosa* samples (Fig. 2B). Moreover, a significant difference among the three cockroach species with respect to microbiome composition was detected using PERMANOVA, which is a non-parametric statistical test for analyzing the differences between the centroids or the dispersion of groups in multivariate datasets (11).

**Figure 2.**
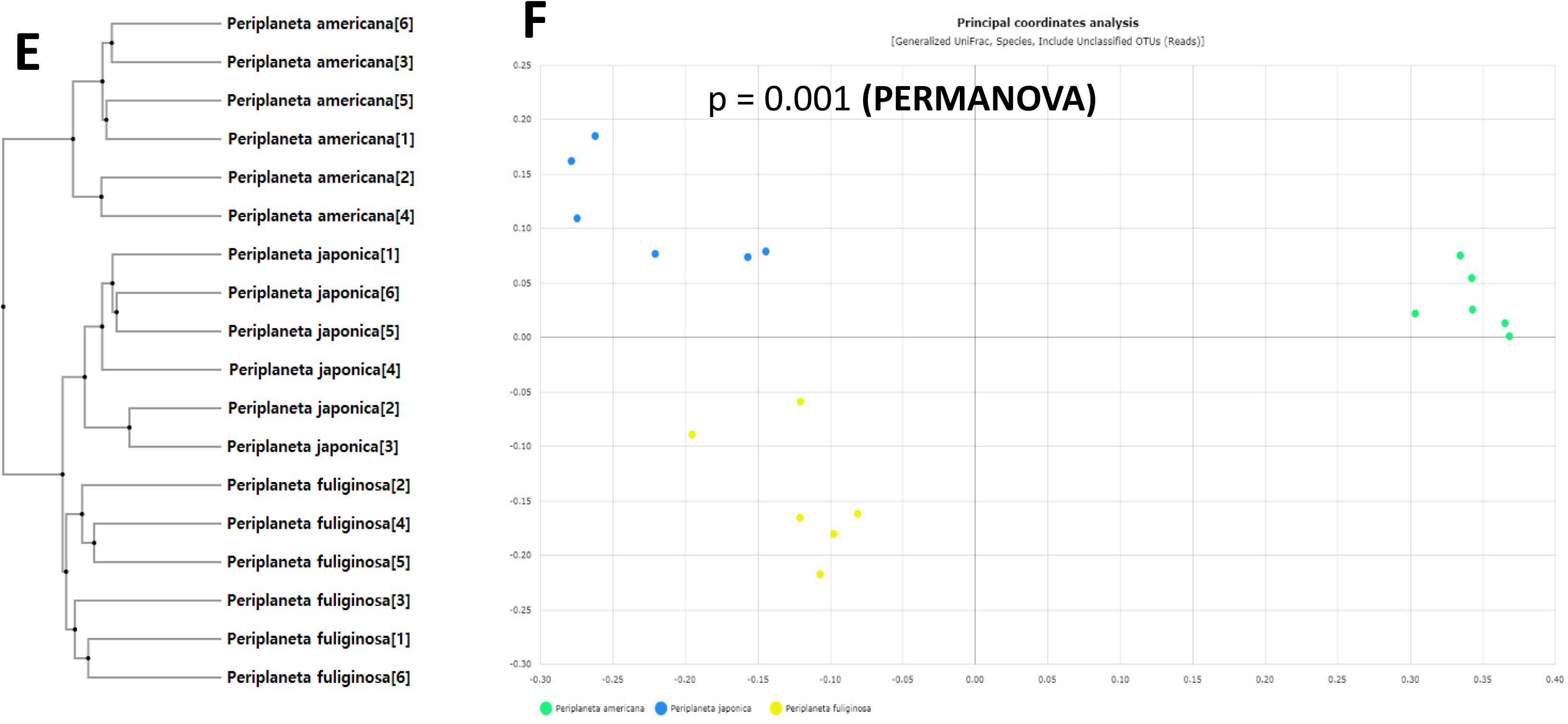
**(A)** Unweighted pair group method with arithmetic mean (UPGMA) clustering and **(B)** principal-coordinate analysis depicting differences in the taxonomic compositions of the bacterial communities between *P. americana* and *P. japonica* and *P. fuliginosa*. *indicates statistically significant differences between the three species of cockroaches (Wilcoxon rank-sum test, p < 0.05).

With respect to the bacterial taxa present in the three cockroach species at the species level (Data Set S2), less than 1% of the bacterial species in *P. americana* accounted for 57.09% of all the microbial species present in *P. americana*. However, bacterial species not included in the aforementioned 1% were more abundant in *P. americana* than in either *P. japonica* or *P. fulginosa*. The endosymbiont *Blattabacterium* CP001429_s accounted for 63.13% and 57.34% of all bacterial species in *P. japonica* and *P. fulginosa*, respectively (Fig. 3A). In *P. japonica*, the endosymbiont *Blattabacterium*_uc, was also present. *Periplaneta japonica* and *P. fulginosa* had many bacterial species in common and had similar compositions of microbial species (Fig. 3A).

**Figure 3.**
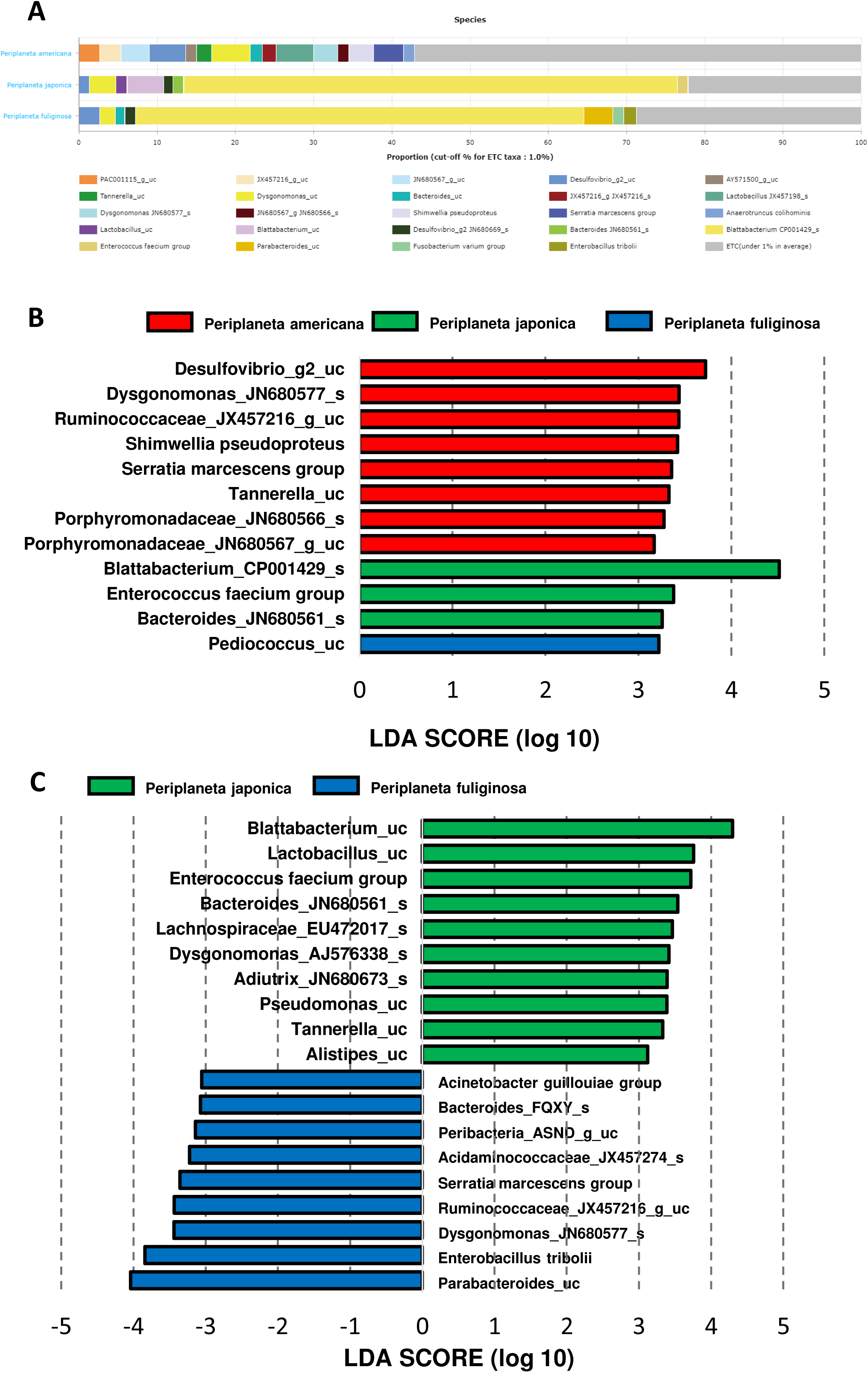
**(A)** The distribution of bacterial taxa at the species level in the cockroach samples in the three species of cockroaches. Each bar depicts the mean relative abundance value of the independent replicates. Bacterial species comprising more than 1% of reads are shown. Each bar depicts the mean relative abundance value of the independent replicates (n = 6, *P. americana*; n = 6, *P. japonica*; n = 6, *P. fuliginosa*). **(B)** LEfSe analysis of differentially abundant bacterial taxa between cockroaches of the three species and **(C)** between *P. japonica* and *P. fuliginosa.* Only taxa meeting a significant LDA threshold (> 3) are shown.

An LEfSe analysis was performed to identify significant differences in bacterial abundance between the cockroach species. The taxa with high LDA scores in *P. americana* were *Desulfovibrio*_g2_uc, *Dysgonomonas*_JN680577_s, and *Serratia marcescens*. In *P. japonica*, *Blattabacterium*_CP001429_s and *Enterococcus faecium* presented high LDA scores. Finally, *Pediococcus*_uc was the species with the highest LDA score in *P. fulginosa* (Fig. 3B). When the bacterial communities in *P. japonica* and *P. fulginosa* were compared without *P. americana*, *Blattabacterium*_uc and *Lactobacillus*_uc were found to be highly abundant in *P. japonica*, while *Parabacteroides*_uc and *Enterobacillus tribolii* were highly abundant in *P. fulginosa* and were the species with the highest LDA scores (Fig. 3C).

## Discussion

Previous studies have shown that microbiomes may differ based on the diets or rearing conditions of their hosts (4). The cockroaches used in this study had lived for many generations under the same conditions, and it was therefore thought that other variables, such as diet, temperature, and humidity would not strongly affect the microbiome.

We evaluated the microbiomes of three cockroach species to determine whether a difference was present among their bacterial profiles. The results indicate that species richness did not differ between cockroach species, but abundance and equity were higher in *P. Americana* than in either *P. japonica* or *P. fuliginosa.*

A previous study reported that the microbiome of *P. americana* was resilient and stable when the cockroach underwent a dietary shift (6). This study found that there were no significant phylum-level differences in the observed microbiomes among the three *P. americana* groups (i.e., diet with six foods, mixed diet, and starvation diet). Furthermore, this study also found no differences between the three *P. americana* groups with respect to either alpha or beta diversity, although differences in microbiome composition that were attributable to environmental factors were observed (6). Similarly, in this study, assuming *P. americana* is stable with regard to dietary shifts, we observed differences in microbial composition attributable to environmental factors at the phylum level in our study (Fig. S2).

With respect to species composition, more than 1% of the species in *P. americana* presented the greatest diversity among the three cockroach species. In *P. japonica* and *P. fuliginosa*, *Blattabacterium* CP001429_s accounted for more than half of all bacterial species. Moreover, another endosymbiont was present in *P. japonica*. In addition, we can see that many bacterial species are present in both microbiomes. Nonetheless, differences between the three cockroach species with respect to the composition of bacterial species were identified using the UPGMA and PCoA clustering analysis. We confirmed that *P. japonica* and *P. fuliginosa* clustered before *P. americana*. In combination with the species composition results, these results suggest that *P. japonica* and *P. fuliginosa* have more similar bacterial compositions compared to that of *P. americana*.

*P. americana* exhibited more prevalent species than either *P. japonica* or *P. fuliginosa*. *Dysgonomonas* species present in *P. americana*, can cause gastroenteritis in immunocompromised individuals (12). *S. marcescens* has been found to be associated with hospital-acquired infections (HAI), and is an opportunistic pathogen that is often involved in urinary tract and wound infections (13). *Blattabacterium_CP001429_s* was present in *P. fuliginosa*, but it was more specific to *P. japonica*. *Enterococcus faecium* can live in the gastrointestinal tract of both humans and animals, but it can cause endocarditis (14). *Periplaneta fuliginosa* had many bacterial species in common with *P. japonica*, but *Pediococcus_uc* was more abundant in all three cockroach species. Nevertheless, PCoA and UPGMA showed different clustering results, and *P. fuliginosa* showed a substantial number of species of bacteria in common with *P. japonica*, although differences were present between the two.

With regard to the limitations of this study, the microbiome composition at the phylum level of *P. americana* did not appear to be affected due to dietary shifts, although it appeared to be affected by environmental factors (6). Nonetheless, the microbiome composition at the phylum level of *P. japonica* and *P. fuliginosa* are unknown and thus it is not possible to conclude that microbiomes would be similar when diets and environmental factors change. In future studies, differences between these two species with regard to diet and environmental shifts should also be established to determine which characteristics the bacteria are determining. Furthermore, this information will be beneficial to identify species-specific cockroach features.

In conclusion, we compared the microbiomes of three *Periplaneta* species and found that there were differences in the bacterial composition of their microbiomes despite being reared under the same conditions for many generations.

## Materials and methods

### Cockroach collection

Three species of cockroaches were collected in the same manner. Rat chow was placed in a glass bottle, and the cockroaches that entered the bottle were collected. These cockroaches were brought back to the laboratory and raised. The *P. americana*, *P. japonica*, and *P. fuliginosa* individuals were collected in Yongsan, Seoul, and Incheon, respectively. *Periplaneta americana* and *P. fuliginosa* were maintained in the laboratory of the Arthropods of Medical Importance Bank of Yonsei University College of Medicine in Seoul, Korea, since 1998, while *P. japonica* was reared since 2017.

### Rearing conditions

The cockroaches used in this study were reared for many generations under the same laboratory conditions to minimize the potential influence of environmental factors and diet. In addition, all cockroaches used in this study were in the adult stage. All cockroaches were reared in plastic boxes (27 × 34 × 19 cm) and incubated at 25 °C. The cockroaches were fed Purina Rat Chow (Basel, Switzerland), containing crude protein, crude fat, crude fiber, crude ash, calcium, and phosphorus) and were supplied tap water *ad libitum*.

### DNA extraction

Each cockroach surface was sterilized using alcohol. The cockroaches were then frozen with liquid nitrogen and individually crushed using a mortar and pestle, and their DNA was extracted. Total DNA was extracted using the NucleoSpin DNA Insect Kit (Macherey-Nagel, Düren, Germany) following the instructions of the manufacturer. Each cockroach sample was separately placed in a bead tube and subjected to the following steps: cell lysis, silica membrane-DNA binding, and silica membrane washing and drying (https://www.mn-net.com/ProductsBioanalysis/DNAandRNApurification/DNA/DNAfromtissueandcells/NucleoSpinDNAInsect/tabid/12727/language/en-US/Default.aspx). The DNA extracted from each sample was eluted in 20 µl of elution buffer. All processing and sequencing procedures were conducted at a clean bench, under a sterilized hood, and in a DNA-free room. DNA concentrations were quantified using a ND-1000 Nanodrop (Thermo-Fisher Scientific, Waltham, MA). The extracted DNA was stored at −80 °C in a deep freezer.

### Amplification of 16S rRNA by polymerase chain reaction (PCR)

The 16S rRNA V3–V4 region was amplified by PCR using forward (5′-TCGTCGGCAGCGTCAGATGTGTATAAGAGACAGCCTACGGGNGGCWGCAG-3′) and reverse primers (5′-GTCTCGTGGGCTCGGAGATGTGTATAAGAGACAGGACTACHVGGGTATCTAATC C-3′) utilizing an Illumina MiSeq V3 cartridge (San Diego, CA) with 600 cycles in accordance with the methodology previously described by Kim et al. (15).

### Next-generation sequencing (NGS)

A limited-cycle amplification step was performed to add multiplexing indices and Illumina sequencing adapters. The libraries were normalized, pooled, and sequenced on the Illumina MiSeq V3 cartridge platform in accordance with the instructions from the manufacturer.

### Bioinformatics and statistics

Bioinformatic analyses were performed following previously described methods (15, 16). Raw reads were processed through a quality check, and low quality (< Q25) reads were removed using Trimmomatic 0.32 (17). Paired-end sequence data were subsequently merged using PandaSeq (18). Primers were then trimmed using the ChunLab in-house program (ChunLab, Inc., Seoul, Korea) by applying a similarity cut-off of 0.8. Sequences were denoised using the Mothur pre-clustering program, which merges sequences, extracts unique sequences, and allows up to two differences between sequences (19). The EzBioCloud database (https://www.ezbiocloud.net/) (16) was used to assign taxonomic information in conjunction with BLAST 2.2.22 (NCBI, Bethesda, MD), and pairwise alignments were generated to calculate similarity (20, 21). The UCHIME algorithm and non-chimeric 16S rRNA database from EzTaxon were used to detect chimeric sequences for reads with a best hit similarity rate < 97% (22). In ChunLab, contigs and singletons that were identified when similarity was < 97% at the taxon-assignment stage were deemed to be chimeras based on the non-chimera database (DB) of the corresponding region. The DB used herein was based on various databases hosted on NCBI and the ChunLab. The bioinformatic ‘usearch’ tool in ChunLab was used to directly remove chimeric reads. Sequence data were then clustered using CD-Hit and UCLUST (23, 24).

All of the described analyses were performed using BIOiPLUG, a commercially available ChunLab bioinformatic cloud platform for microbiome research (https://www.bioiplug.com/). Rarefaction for the obtained OTUs was calculated using the ChunLab pipeline, in accordance with the methodology of Heck et al. (25). The reads were normalized to 43,000 to perform the analyses. We computed the Shannon index (26) and performed unweighted pair group method with arithmetic mean (UPGMA) clustering (27), principal coordinates analysis (PCoA) (28), and a permutational multivariate analysis of variance (PERMANOVA) (29) based on the generalized UniFrac distance (30). We used the Wilcoxon rank-sum test to evaluate the differences in the number of OTUs and used the Shannon, phylogenetic, and Pielou indexes to compare microbiome diversity between the three cockroach species. We used linear discriminant analysis (LDA) effect size (LEfSe) analysis to identify significantly different taxa between the two and three species of cockroaches (31).

## Data availability

This data is underway (Raw data for NGS).

## Acknowledgements

This study was supported by a National Research Foundation of Korea (NRF) grant funded by the Korean Government (MEST; numbers NRF-2019R1A2B5B01069843).

## Supplemental Material

**Figure. S1** Rarefaction curves of the number of operational taxonomic units (OTUs) of the three groups of cockroaches. **(A)** Rarefaction curves of *Periplaneta americana*. **(B)** Rarefaction curves of *P. japonica*. **(C)** Rarefaction curves of *P. fuliginosa*.

**Figure. S2** The distribution of bacterial taxa at the phylum level in the cockroach samples from the three species. Each bar depicts the mean relative abundance value of the independent replicates. Bacterial phyla comprising more than 1% of the reads are shown. Each bar depicts the mean relative abundance value of the independent replicates (n = 6, *P. americana*; n = 6, *P. japonica*; n = 6, *P. fuliginosa*).

**Data Set S1** Alpha diversity numerical value data in the three cockroach species.

**Data Set S2** List of taxa found in the three cockroach species.

## References

1. Appel AG. 1991. Water relations and thermal sensitivity of several cockroach species (Dictyoptera: Blattidae and blaberidae). Comp Biochem Physiol 100:353–356.

2. Patourel GNJ. 1993. Cold-tolerance of the oriental cockroach *Blatta orientalis*. Entomol Exp Appl 68:257–263.

3. Mullins DE. 2015. Physiology of Environmental Adaptations and Resource Acquisition in Cockroaches. Annu Rev Entomol 60:473–492.

4. Vicente CSL, Mondal SI, Akter A, Ozawa S, Kikuchi T, Hasegawa K. 2018. Genome analysis of new *Blattabacterium* spp., obligatory endosymbionts of *Periplaneta fuliginosa* and *P. japonica*. PLoS One 13:e0200512.

5. Kakumanu ML, Maritz JM, Carlton JM, Schal C. 2018. Overlapping Community Compositions of Gut and Fecal Microbiomes in Lab-Reared and Field-Collected German Cockroaches. Appl Environ Microbiol 84:e01037–18.

6. Tinker KA, Ottesen EA. 2016. The Core Gut Microbiome of the American Cockroach, *Periplaneta americana*, Is Stable and Resilient to Dietary Shifts. Appl Environ Microbiol 82:6603–6610.

7. Roth LM. 1981. Introduction, p 1–14. In Bell WJ, Adiyodi KG (ed), The American Cockroach. Springer, New York, NY.

8. Rust MK, Reierson DA, Hansgen KH. 1991. Control of American cockroaches (Dictyoptera: Blattidae) in sewers. J Med Entomol 28:210–213.

9. Jeong KY, Hwang H, Lee J, Lee IY, Kim DS, Hong CS, Ree HI, Yong TS. 2004. Allergenic characterization of tropomyosin from the dusky brown cockroach, *Periplaneta fuliginosa*. Clin Diagn Lab Immunol 11:680–685.

10. Tanaka K, Tanaka S. 1997. Winter Survival and Freeze Tolerance in a Northern Cockroach, *Periplaneta japonica* (Blattidae: Dictyoptera). Zoolog Sci 14:849–853.

11. Ericsson AC, Gagliardi J, Bouhan D, Spollen WG, Givan SA, Franklin CL. 2018. The influence of caging, bedding, and diet on the composition of the microbiota in different regions of the mouse gut. Sci Rep 8:4065.

12. Jayasudha R, Chakravarthy SK, Prashanthi GS, Sharma S, Garg P, Murthy SI, Shivaji S. 2018. Alterations in gut bacterial and fungal microbiomes are associated with bacterial Keratitis, an inflammatory disease of the human eye. J Biosci 43:835–856.

13. Khanna A, Khanna M, Aggarwal A. 2013. *Serratia marcescens*-a rare opportunistic nosocomial pathogen and measures to limit its spread in hospitalized patients. J Clin Diagn Res 7:243–246.

14. Munita JM, Arias CA, Murray, BE. 2012. Enterococcal endocarditis: can we win the war? Curr Infect Dis Rep 14:339–349.

15. Kim JY, Yi MH, Hwang Y, Lee JY, Lee IY, Yong D, Yong TS. 2018. 16S rRNA profiling of the *Dermatophagoides farinae* core microbiome: *Enterococcus* and *Bartonella*. Clin Exp Allergy 48:607–610.

16. Yoon SH, Ha SM, Kwon S, Lim J, Kim Y, Seo H, Chun J. 2017. Introducing EzBioCloud: a taxonomically united database of 16S rRNA gene sequences and whole-genome assemblies. Int J Syst Evol Microbiol 67:1613–1617.

17. Bolger AM, Lohse M, Usadel B. 2014. Trimmomatic: a flexible trimmer for Illumina sequence data. Bioinformatics 30:2114–2120.

18. Masella AP, Bartram AK, Truszkowski JM, Brown DG, Neufeld JD. 2012. PANDAseq: paired-end assembler for illumina sequences. BMC Bioinformatics 13:31.

19. Schloss PD, Westcott SL, Ryabin T, Hall JR, Hartmann M, Hollister EB, Lesniewski RA, Oakley BB, Parks DH, Robinson CJ, Sahl JW, Stres B, Thallinger GG, Van Horn DJ, Weber CF. 2009. Introducing mothur: open-source, platform-independent, community supported software for describing and comparing microbial communities. Appl Environ Microbiol 75:7537–7541.

20. Myers EW, Miller W. 1988. Optimal alignments in linear space. Comput Appl Biosci 4:11–17.

21. Altschul SF, Gish W, Miller W, Myers EW, Lipman DJ. 1990. Basic local alignment search tool. J Mol Biol 215:403–410.

22. Edgar RC, Haas BJ, Clemente JC, Quince C, Knight R. 2011. UCHIME improves sensitivity and speed of chimera detection. Bioinformatics 27:2194–2200.

23. Edgar RC. 2010. Search and clustering orders of magnitude faster than BLAST. Bioinformatics 26:2460–2461.

24. Fu L, Niu B, Zhu Z, Wu S, Li W. 2012. CD-HIT: accelerated for clustering the next-generation sequencing data. Bioinformatics 28:3150–3152.

25. Heck KL, van Belle G, Simberloff D. 1975. Explicit calculation of the rarefaction diversity measurement and the determination of sufficient sample size. Ecology 56:1459–1461.

26. Shannon C, Petigara N, Seshasai S. 1948. A mathematical theory of communication. Bell Syst Tech J 27:379–423.

27. Sneath PHA, Sokal RR (2nd ed). 1973. p 573, Numerical Taxonomy: The Principles and Practice of Numerical Classification. WH Freeman and Co., San Francisco, CA

28. Gower JC. 1966. Some distance properties of latent root and vector methods used in multivariate analysis. Biometrika 53:325–338.

29. Anderson MJ. 2001. A new method for non-parametric multivariate analysis of variance. Austral Eco 26:32–46.

30. Lozupone C, Knight R. 2005. UniFrac: a new phylogenetic method for comparing microbial communities. Appl Environ Microbiol 71:8228–8235.

31. Segata N, Izard J, Waldron L, Gevers D, Miropolsky L, Garrett WS, Huttenhower C. 2011. Metagenomic biomarker discovery and explanation. Genome Biol 12:R60.

